# Loss of the APP regulator RHBDL4 preserves memory in an Alzheimer’s disease mouse model

**DOI:** 10.1101/2024.02.22.579698

**Authors:** Ylauna Christine Megane Penalva, Sandra Paschkowsky, Jingyun Yang, Sherilyn Junelle Recinto, Jessica Cinkorpumin, Marina Ruelas Hernandez, Bin Xiao, Albert Nitu, Helen Yee-Li Wu, Hans Markus Munter, Bernadeta Michalski, Margaret Fahnestock, William Pastor, David A. Bennett, Lisa Marie Munter

## Abstract

Characteristic cerebral pathological changes of Alzheimer’s disease (AD) such as glucose hypometabolism or the accumulation of cleavage products of the amyloid precursor protein (APP), known as Aβ peptides, lead to sustained endoplasmic reticulum (ER) stress and neurodegeneration. To preserve ER homeostasis, cells activate their unfolded protein response (UPR). The rhomboid-like-protease 4 (RHBDL4) is an enzyme that participates in the UPR by targeting proteins for proteasomal degradation. We demonstrated previously that RHBLD4 cleaves APP in HEK293T cells, leading to decreased total APP and Aβ. More recently, we showed that RHBDL4 processes APP in mouse primary mixed cortical cultures as well. Here, we aim to examine the physiological relevance of RHBDL4 in the brain. We first found that brain samples from AD patients and an AD mouse model (APPtg) showed increased RHBDL4 mRNA and protein expression. To determine the effects of RHBDL4’s absence on APP physiology *in vivo*, we crossed APPtg mice to a RHBDL4 knockout (R4^-/-^) model. RHBDL4 deficiency in APPtg mice led to increased total cerebral APP and amyloidogenic processing when compared to APPtg controls. Contrary to expectations, as assessed by cognitive tests, RHBDL4 absence rescued cognition in 5-month-old female APPtg mice. Informed by unbiased RNAseq data, we demonstrated *in vitro* and *in vivo* that RHBDL4 absence leads to greater levels of active β-catenin due to decreased proteasomal clearance. Decreased β-catenin activity is known to underlie cognitive defects in APPtg mice and AD. Our work suggests that RHBDL4’s increased expression in AD, in addition to regulating APP levels, leads to aberrant degradation of β-catenin, contributing to cognitive impairment.

## Introduction

Alzheimer’s disease (AD) is a debilitating neurodegenerative disorder characterized by progressive impairments in cognitive functions and memory loss (1). It accounts for 50% to 60% of all dementia cases, with dementia being recognized as the 5^th^ leading cause of death worldwide (2, 3). While therapeutic strategies are actively undergoing development, continued clinical trial failures suggest that a cure or effective treatments are still far from reach (4, 5). The difficult drug discovery process may be due to the complexity of the disease, as multiple factors such as cerebrovascular dysregulation (6, 7), abnormal glucose metabolism (8, 9), and aggregates of misfolded proteins (10–12) have been involved in AD onset and progression. One of the most extensively studied pathological hallmarks is the accumulation of Aβ peptides as extracellular plaques in the brain (13, 14). Aβ peptides stem from the sequential proteolytic cleavage by β-secretase and γ-secretase of the amyloid precursor protein (APP) (15), a type-I transmembrane glycoprotein which is genetically causative for AD through various inherited mutations (16, 17). While APP’s function is linked in part to synaptogenesis, axonal outgrowth, and myelination, it is still unclear how APP’s dysfunction may lead to AD pathogenesis (18–20).

To approach this question, it is imperative to better understand APP regulation in the cell and in the brain. We have described a novel processing pathway of APP mediated by the rhomboid-like-protease 4 (RHBDL4) where APP is cleaved by RHBDL4 at multiple sites in the endoplasmic reticulum (ER) (21, 22). We showed in HEK293T cells that RHBDL4 cleavage of APP led to the production of RHBDL4-specific APP fragments and, decreased cell surface levels of APP and Aβ production (21). We further confirmed that RHBDL4 cleavage of APP occurs in primary mixed cortical cultures (23). Specifically, we found that RHBDL4 knockout (R4^-/-^) mixed cortical cultures did not produce an RHBDL4-specific APP fragment termed “A_η_-like” for its resemblance to the APP fragment originally described to be produced by MT5-MPP by Willem et al. (23, 24). We also observed that APP expression was elevated in brain lysates from R4^-/-^ mice in comparison to controls, suggesting a physiological relevance of RHBDL4 for APP *in vivo* (23).

It is important to note that RHBDL4 has been described by other groups as an important player in the unfolded protein response (UPR). In particular, it participates in ER associated degradation (ERAD) pathways (25, 26). These pathways are activated to protect ER homeostasis when cells undergo ER stress (27). RHBDL4 can recognize ubiquitinated transmembrane proteins via its ubiquitin interacting motif (UIM), cleave them and later target them for proteasomal degradation through VCP/p97 mediated extraction from the membrane (25). More recently, it was shown that RHBDL4 also targets aggregated proteins that do not have a transmembrane domain for degradation through the proteasome via interactions with partners, Erlin1/2, that bring aggregated targets directly to RHBDL4 (28). The diversity of RHBDL4’s substrates and interactors suggests it plays a major role in maintaining ER homeostasis (29–31). In the context of AD, the UPR is upregulated due to ER stress induced by oxidative stress, metabolic restrictions, and misfolded protein accumulation (32, 33). Given our and others’ findings, we were foremost interested in determining RHBDL4’s potential involvement in AD pathology and behaviour.

In the present study, we assessed RHBDL4 expression at the mRNA and protein level in AD patients and in adult APP transgenic J20 (APPtg) mouse brain samples. We further explored APP expression and amyloidogenic processing in the absence of RHBDL4 in the APPtg model. We also evaluated the effects of RHBDL4 deficiency on cognitive performance in the APPtg model, which led us to explore β-catenin expression and activity *in vitro* and *in vivo* in the R4^-/-^ model.

## Results

### RHBDL4 mRNA and protein expression levels are increased in brain samples from AD subjects and APPtg mice as compared to controls

First, we analysed the expression pattern of RHBDL4 in AD dementia cases compared to those without dementia using samples from the Religious Orders Study or Rush Memory and Aging Project (ROSMAP) cohorts. AD diagnosis was performed according to NIA Reagan Pathology criteria (34). RNA-sequencing analysis revealed higher expression of a transcript encoding functional, full length RHBDL4 (ENST00000392062.2, RHBDL4 gene name is *RHBDD1*) in subjects with AD dementia compared to non-demented controls (0.24 vs. 0.18 log2 counts per million (cpm), p=0.006; Fig. 1A). Multivariable logistic regression controlling for age at death, sex and education also indicated that RHBDL4 mRNA expression was associated with increased odds of pathologic AD (odds ratio (OR)=2.38, 95% confidence interval (CI): 1.19-4.94; p=0.017) (Table 1). Moreover, in multiple linear regression analyses controlling for age at death, sex and education, we also found that this transcript was negatively associated with global cognition (Fig. 1B). Further analysis indicated that its association with neurofibrillary tangles was mediated by β-amyloid (Table 1). We did not find a significant association of the expression of the transcript with macro- or micro-infarcts, or Lewy bodies (data not shown). Analysis of RHBDL4 protein levels in brain samples from AD cases compared to control samples revealed a trend towards increased levels in AD (Fig. 1C). To explore whether this relationship was conserved in AD mouse models, we assessed RHBDL4 expression in whole brain lysates of 5-month-old APP transgenic J20 (APPtg) mice (35). APPtg mice overexpress human APP with two mutations, Swedish and Indiana, linked to familial Alzheimer’s disease (35). Our results showed a significant increase of both mRNA and protein expression of RHBDL4 in APPtg brains in comparison to wild type (Fig. 1D-E). These conserved changes between humans and mouse model implicate RHBDL4 in the response to or etiology of AD.

**Figure 1:**
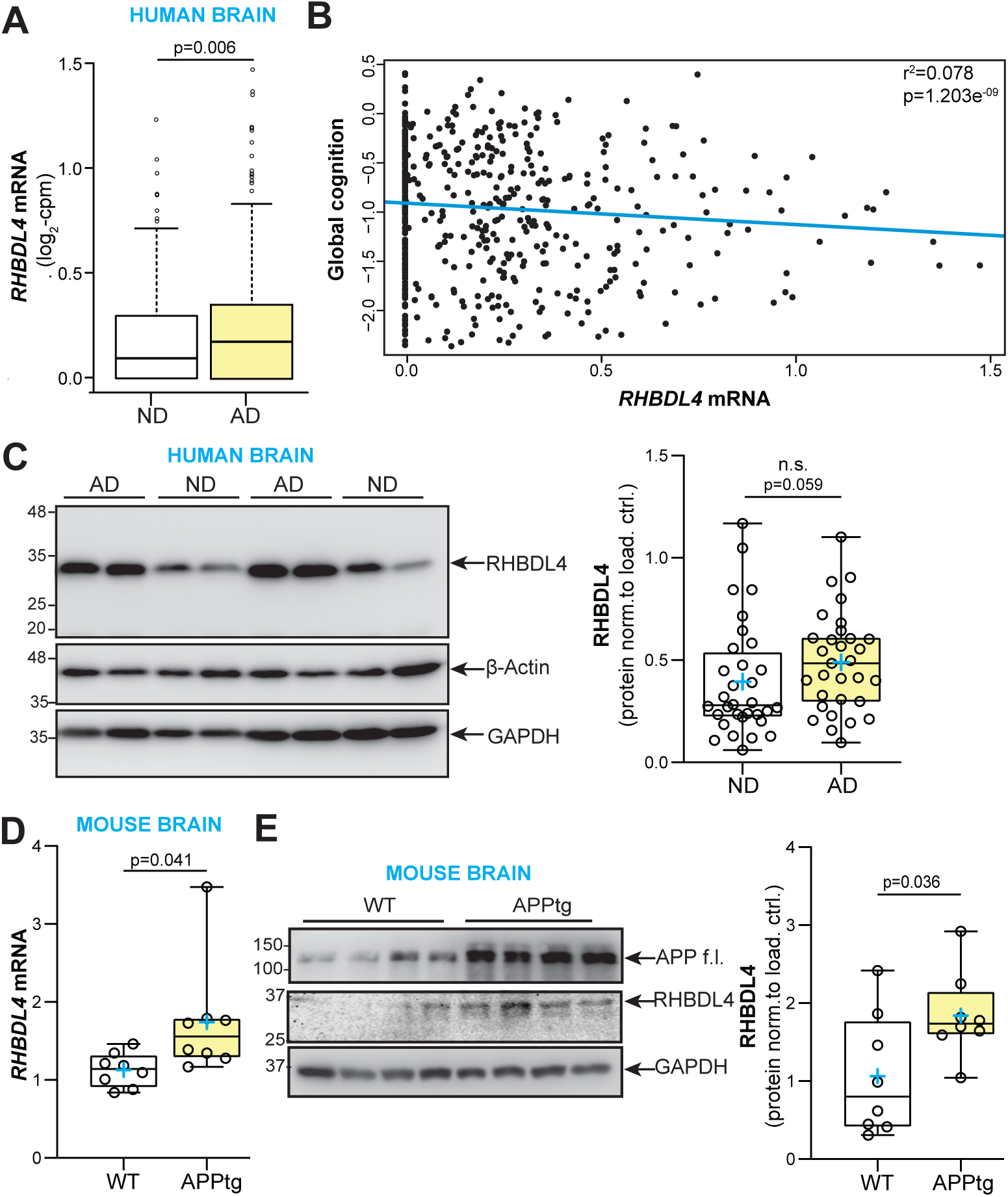
RHBDL4 expression is increased in AD dementia and APPtg mouse brains. A. Analysis of RNA-sequencing data from the Religious Order Study cohort (n=321 AD dementia cases and n=220 no dementia cases). Shown are expression levels of the RHBDL4 mRNA ENST00000392062.2 (one of two transcripts encoding for RHBDL4 full length protein). ND= No dementia samples, AD= AD dementia samples. The line within each box indicates the median of the dataset. The lower and upper edges of the box represent the first (Q1) and third (Q3) quartiles, respectively. The ‘whiskers’ extend from the quartiles to the last data point within 1.5 times the interquartile range (IQR) from the quartile. Data points beyond this range are considered outliers and are plotted individually. B. Scatterplot for association of global cognition and RHBDL4 mRNA (ENST00000392062.2) levels. Results are derived from multiple linear regression analysis controlling for age at death, sex and education; Data were transformed using Box-Cox with negatives. The blue line represents estimated global cognition (transformed using Box-Cox with negatives) for a female with mean level of education (16.5 years) and mean age at death (88.4 years). Association statistics: adjusted r^2^: 0.078, F-statistic: 12.39 on 4 and 535 DF, p-value: 1.203e-09. C. RHBDL4 protein in AD patient brain samples compared to controls (no dementia, no pathology). RHBDL4 protein expression was quantified and normalized to β-actin+GAPDH. n= 31-32 per group, box and whisker plots represent minimum to maximum values with median center lines while blue “+” represents the mean; Mann-Whitney U test after significant Shapiro Wilk test for normality. D. RHBDL4 mRNA expression in APPtg brain samples compared to WT. RHBDL4 mRNA expression normalized to reference genes RSP18 and GAPDH. n=8 per group, box and whisker plots represent minimum to maximum values with median center lines while blue “+” represents the mean; Two-tailed, unpaired t-test performed, and p-value reported. E. RHBDL4 protein expression in APPtg brain samples compared to WT. APP full length (f.l.) is blotted to show overexpression in APPtg model. Low abundance RHBDL4 protein expression was quantified and normalized to GAPDH. RHBDL4 was detected via fluorescence imaging. APP and GAPDH were detected with chemiluminescence imaging. n= 8 per group, box and whisker plots represent minimum to maximum values with median center lines while blue “+” represents the mean; Two-tailed, unpaired t-test performed, and p-value reported.

**Table 1:** Results for linear regression analysis with AD pathology as dependent variable, controlled for age at death, sex, and education, SE: standard error.

| ENST00000392062.2.<br>estimated coefficient | SE | t-stat | p-value | AD response |
| --- | --- | --- | --- | --- |
| 0.547 | 0.170 | 3.215 | 0.0014 | β-amyloid |
| 2.601 | 1.224 | 2.125 | 0.0340 | tangles |
| 0.284 | 0.094 | 3.019 | 0.0027 | AD pathology |

### Deletion of RHBDL4 in the APPtg mouse model affects APP protein levels and APP amyloidogenic processing in the brain

To determine the effects of RHBDL4 on APP homeostasis *in vivo* in the context of a preclinical mouse model of AD, we crossed the R4^-/-^ strain (30) to the APPtg model (35). Our breeding scheme was designed to yield wild type (WT), APPtg, R4^-/-^, APPtg/R4^-/+^, and APPtg/R4^-/-^ littermates (Fig. 2A). Mice were tested at 5 months of age for cognitive functions and biochemical assays were performed on collected organs to gain mechanistic insight (Fig. 2A). We chose this age for our tests as it is a pivotal time point where APPtg mice start to show cognitive impairment and early β-amyloid plaque deposition in comparison to WT mice (35, 36). We first confirmed the expected absence of RHBDL4 protein in brain pancreas and liver tissues (Fig. 2B). Acknowledging that RHBDL4 expression is high in liver, we confirmed the RHBDL4 genotype of all experimental groups on liver lysates (Fig. 2C). Thereafter, we examined total APP expression in the brains of our mice. First, we confirmed our findings that R4^-/-^ mice show increased endogenous APP levels as compared to WT (23). In addition, in the APPtg model, RHBDL4 loss resulted in higher total APP protein expression (Fig. 2D). These findings corroborate the idea that RHBDL4 contributes to the regulation of APP levels and potentially its availability for physiological functions. To address this point, we additionally looked at markers of amyloidogenic processing which we had previously found to be affected by RHBDL4 overexpression in cell culture (21). First, we found that β-secretase-derived C-terminal fragments (β-CTFs) of APP were significantly increased in APPtg/R4^-/+^ as compared to APPtg mice (Fig. 2D). In addition, we quantified Aβ levels which are a typical biomarker of AD and are routinely studied to assess progression of the disease. ELISA measurements of DEA soluble Aβ_38_, Aβ_40_ and Aβ_42_ from the brain showed that Aβ_40_ levels rose significantly when RHBDL4 was knocked out in female APPtg, with a trend towards an increase for the other Aβ species (Fig 2E). We did not observe any significant effect of RHBDL4 knockout on the Aβ_42_/ Aβ_40_ or Aβ_42_/total Aβ_38+40+42_ ratios (Fig 2F). Formic acid soluble plaque-derived Aβ levels were low at 5 months of age, as expected, since plaque load is low at this early time point (35). However, we detected formic acid soluble Aβ_42_, which showed a trend towards an increase in APPtg mice in the absence of RHBDL4 (Fig. 2G). Altogether, the absence of the RHBDL4-mediated processing pathway may increase the availability of APP for amyloidogenic processing.

**Figure 2:**
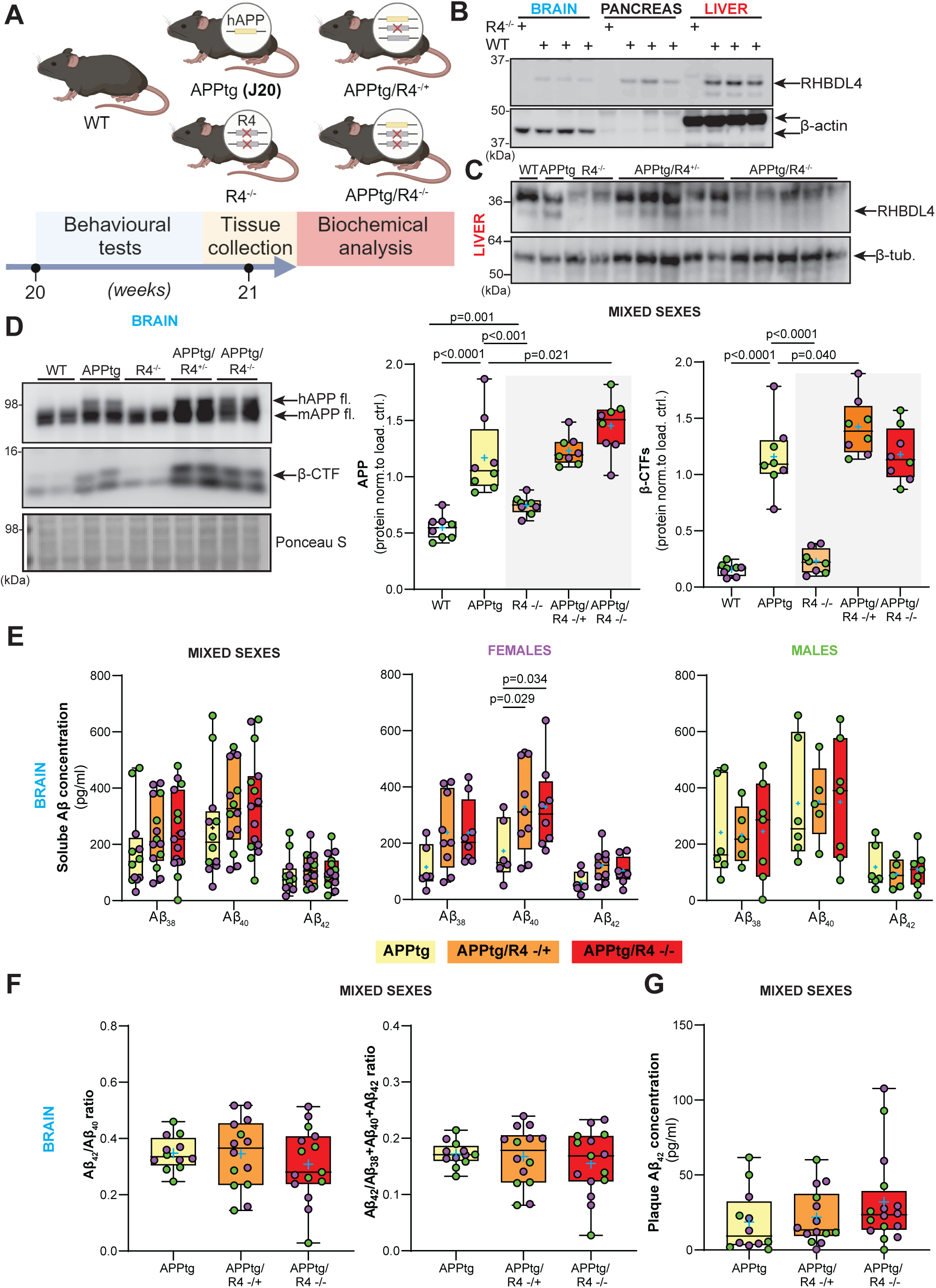
Depletion of RHBDL4 expression in APPtg mice brains leads to higher APP production and amyloidogenic processing. A. Schematic representation of experimental design and analysis timeline for APPtg J20 mice crossed to the R4^-/-^ model. B. RHBDL4 expression in brain, pancreas and liver lysates of WT mice. Lysates from RHBDL4 KO brains are used as a negative control for RHBDL4 immunoblot. β-actin as loading control. C. RHBDL4 expression in liver lysates of WT, APPtg, R4^-/-^, APPtg/R4^-/+^, and APPtg/R4^-/-^ mice. β-tubulin as loading control. D. Total (mutant and endogenous) APP and β-CTFs expression in brain lysates of WT, APPtg, R4^-/-^, APPtg/R4^-/+^, and APPtg/R4^-/-^ mice. Expression was quantified and normalized to Ponceau-S. n= 8 per group, box and whisker plots represent minimum to maximum values with median center lines while blue “+” represents the mean; For APP: Two-tailed, unpaired t-test performed between WT and R4^-/-^ group and one-way ANOVA (p<0.0001) with Holm-Sidak multiple comparison test where APPtg is compared to all other groups. For β-CTFs: one-way ANOVA (p<0.0001) with Holm-Sidak multiple comparison test where APPtg is compared to all other groups. Significant p-values for t-test and post hoc analysis reported. E. DEA soluble Aβ38, Aβ40, and Aβ42 concentration in APPtg, APPtg/R4^-/+^, and APPtg/R4^-/-^ brains. n= 12-15 for mixed sexes, n=6-9 for females (purple) and n=5-7 for males (green). Box and whisker plots represent minimum to maximum values with median center lines while blue “+” represents the mean; For mixed sexes: two-way ANOVA (non-significant interaction, p<0.0001 for Aβ species main effect and non-significant genotype main effect) with Tukey’s multiple comparison test. For females: two-way ANOVA (non-significant interaction, p<0.0001 for Aβ species main effect and p=0.0032 for the genotype main effect) with Tukey’s multiple comparison test. For males: two-way ANOVA (non-significant interaction, p=0.0001 for Aβ species main effect and non-significant genotype main effect) with Tukey’s multiple comparison test. Significant p-values for post hoc analysis reported. F. Aβ42/Aβ40 or Aβ42/total Aβ38+40+42 ratios (DEA soluble) in APPtg, APPtg/R4^-/+^, and APPtg/R4^-/-^ brains. n=12-15 per group, mean ± SEM; non-significant one-way ANOVA (p=0.578 for Aβ42/ Aβ40 ratio and p= 0.635 for Aβ42/total Aβ38+40+42 ratio). G. Formic acid soluble Aβ42 concentration in APPtg, APPtg/R4^-/+^, and APPtg/R4^-/-^ brains. n=12-15 per group, mean ± SEM; non-significant one-way ANOVA (p= 0.304) D-G. Female data points are in purple and male in green.

### Cognition is preserved in APPtg female mice in the absence of RHBDL4 expression

As the RHBDL4 knockout positively regulates the expression of APP and the production of amyloidogenic fragments in APPtg brains, we were curious to assess RHBDL4’s role as a potential early modulator of cognitive behaviour. As expected, in the Y maze, APPtg mice performed worse than WT mice as shown by a lower spontaneous alternation performance (SAP) score (Fig. 3A). We also found that R4^-/-^ mice did not show any memory defect with this test (Fig. 3A). On the other hand, we did not observe a significant difference in performance between the APPtg and the APPtg/R4^-/+^ or APPtg/R4^-/-^ mice (Fig. 3A). Therefore, we further stratified our data by sex as it has been described that sex impacts onset and severity of cognitive decline in AD mouse models, with female mice generally being impacted earlier than their male counterparts (37). While female APPtg mice performed worse than WT and R4^-/-^ in the Y maze test, males did not show similar memory deficits yet with this assessment, which may be attributed to the young age of the mice (Fig. 3A). Furthermore, female APPtg mice exhibited worse performance than the APPtg/R4^-/+^ or APPtg/R4^-/-^ groups, suggesting that altering RHBDL4 expression is sufficient to rescue cognitive impairments early in disease (Fig. 3A). To confirm these findings, the same groups of mice also underwent the novel object recognition (NOR) test. The unstratified data from the testing phase of the NOR did not reveal any significant relationship between the 5 groups, but there was a trend towards poorer learning performance by the APPtg mice in comparison to WT and R4^-/-^ mice as indicated by the index of preference score (Fig. 3B). Once stratified by sex, we obtained similar results to those of the Y maze. Female APPtg showed learning memory defects in comparison to WT or R4^-/-^ and those defects were rescued in the APPtg/R4^-/+^ or APPtg/R4^-/-^ genotypes (Fig. 3B). Male APPtg mice on the other hand did not behave differently from WT at that age (Fig. 3B). We conclude that at the young age of 5 months, the absence of RHBDL4 in APPtg mice preserved memory in female mice. These findings were overall supported in the McGill-Thy1-APP transgenic mouse model (38) where the absence of RHBDL4 also led to memory preservation during the Y-maze test (Supplemental Fig. 1A, B).

**Figure 3:**
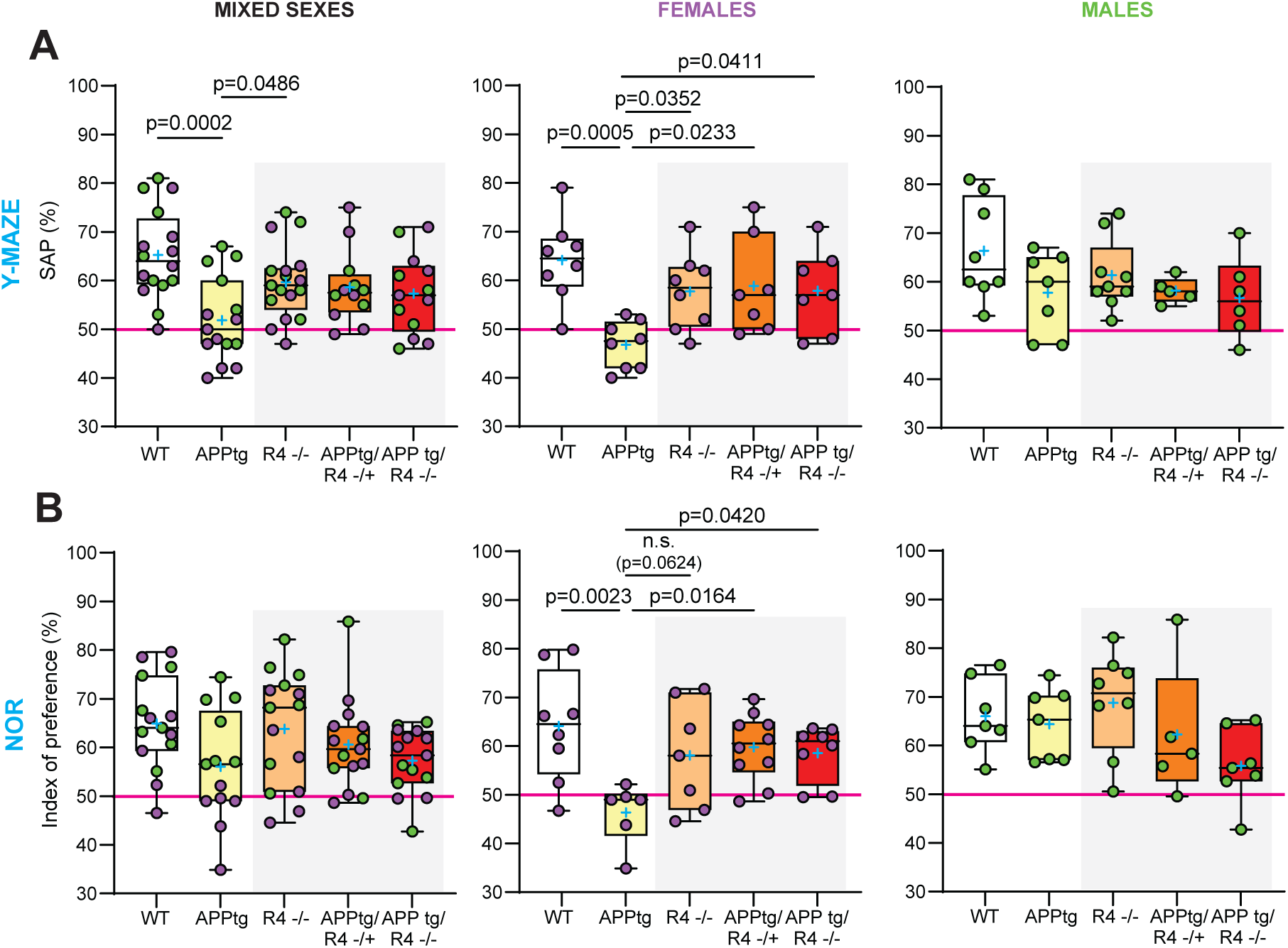
Cognitive defects are improved in the absence of RHBDL4 expression in female APPtg mice. A. Spontaneous alternation performance (SAP) score from Y maze test of WT, APPtg, R4^-/-^, APPtg/R4^-/+^, and APPtg/R4^-/-^ mice. n= 12-17 per group for mixed sexes, n=7-8 per group for females (purple) and n=5-9 per group for males (green). Box and whisker plots represent minimum to maximum values with median center lines while blue “+” represents the mean. One-way ANOVA (p=0.002 for mixed sexes, p= 0.003 for females and p= 0.172 for males) with Dunnett’s multiple comparison test for significant ANOVAs, significant p-values for post hoc analysis reported. B. Index of preference score from novel object recognition (NOR) test of WT, APPtg, R4^-/-^, APPtg/R4^-/+^, and APPtg/R4^-/-^ mice. n= 13-15 per group for mixed sexes, n=6-10 per group for females (purple) and n=5-8 per group for males (green). Box and whisker plots represent minimum to maximum values with median center lines while blue “+” represents the mean. One-way ANOVA (p=0.077 for mixed sexes, p= 0.012 for females and p= 0.129 for males) with Dunnett’s multiple comparison test for significant ANOVAs, significant p-values for post hoc analysis reported.

### β-catenin expression and activity are increased due to defective proteasomal degradation in the absence of RHBDL4 *in vitro*

To discern how RHBDL4 ablation led to a better performance in cognitive tests for impaired female APPtg mice, we aimed to establish which major signalling pathways may be influenced by RHBDL4 loss. Since mouse embryonic fibroblasts (MEFs) express RHBDL4 well (Fig. 4C) and our group had access to R4^-/-^ MEFs, we chose to use them as a simple *in vitro* model for follow-up experiments. First, we collected RNA from WT and R4^-/-^ MEFs and generated cDNA libraries for RNAseq. Using our dataset, we performed differential gene expression analyses and applied p-value cut-off of 0.05 and a fold change cut-off of 1.8. We found that 1304 genes were upregulated while 537 genes were downregulated in R4^-/-^ in comparison to WT MEFs (Fig. 4A). When we completed a KEGG pathway analysis with the upregulated genes using the Enrichr enrichment analysis tool, we observed that the most significantly upregulated pathway in R4^-/-^ MEFs was the Wnt/β-catenin signalling pathway (Fig. 4B). Specifically, 25 upregulated Wnt/β-catenin signalling genes were identified in the absence of RHBDL4, mostly Wnt ligands and receptors for those ligands (Figure 4B, C). We assessed protein expression of one of the upregulated receptors, LRP6, a potent signal transducer of the Wnt/β-catenin signalling pathway (39). We found a striking upregulation at the protein level which corroborated our RNAseq data. This overall piqued our interest as stimulation of this pathway has been established as a possible therapeutic strategy for AD due to its involvement in synapse maintenance and formation as well as axonal pathfinding and dendritogenesis (40, 41). In particular, reduced β-catenin levels, as well as Wnt/β-catenin signalling, have been observed in the brains of AD patients (42, 43) and promotion of β-catenin expression and activity has been shown to rescue cognition in the J20 APPtg model (44–46).

**Figure 4:**
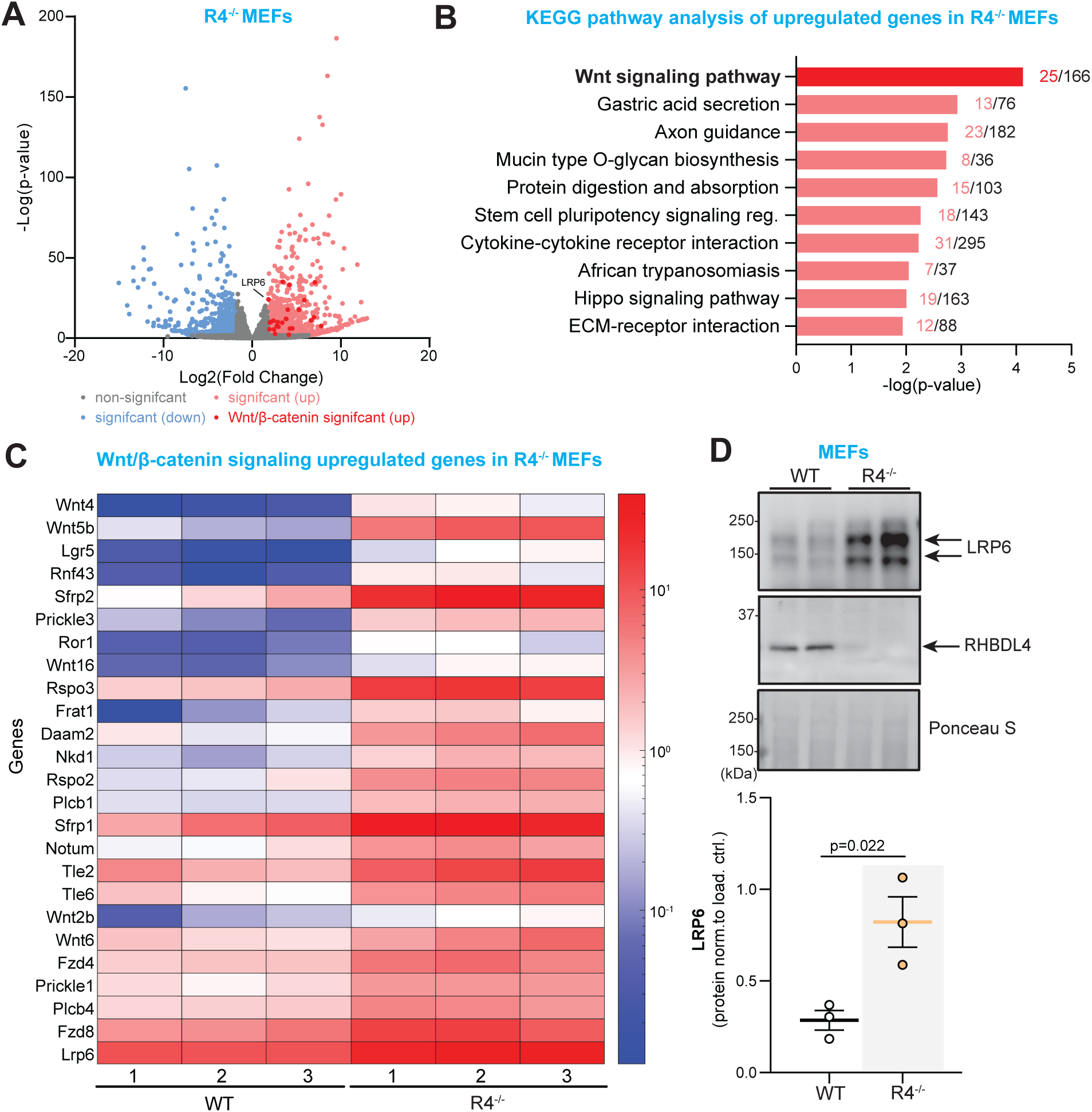
RNAseq data from RHBDL4 null MEFs reveals upregulation of Wnt signalling. A. Volcano plot illustrating differentially expressed genes within R4^-/-^ MEFs relative to wildtype counterparts. Threshold for differential expression was set to log2-fold change of 1.8 and statistical significance set at p-value < 0.05. Genes below those thresholds are in grey. Significantly downregulated genes in R4^-/-^ are represented in light blue while significantly upregulated genes are represented in light red. Wnt signalling upregulated genes are in dark red. B. Illustration adapted from Enrichr of top 10 significantly upregulated KEGG pathways in R4^-/-^ MEFs as compared to control are shown. p-value is computed from the Fisher exact test on Enrichr. C. Heatmap of upregulated Wnt signalling genes in R4^-/-^ MEFs showing normalized expression levels across biological replicates for both R4^-/-^ and WT controls. Each column represents a biological replicate for both genotypes and each row is a gene. D. LRP6 expression in R4^-/-^ and WT MEFs. Expression was quantified and normalized to Ponceau-S. n= 3, mean ± SEM; Two-tailed, unpaired t-test, p-values reported.

Therefore, we assessed β-catenin expression and activity in R4^-/-^ MEFs to further our RNAseq findings. We found that total β-catenin levels were 2-fold increased in R4^-/-^ MEFs in comparison to WT (Fig. 5A). In addition, we analyzed levels of phosphorylated β-catenin (pβ-catenin) at inhibitory positions serine 33, serine 37 and threonine 41. These phosphorylations are commonly used as an indicator of β-catenin inactivity since they render β-catenin transcriptionally inactive and target it for proteasomal degradation (47, 48). No difference in pβ-catenin levels was detected between R4^-/-^ and WT MEFs. Thus, the increased total β-catenin levels in R4^-/-^ MEFs, reflected in their lower pβ-catenin/β-catenin ratios (Fig 5A), suggests that β-catenin function is increased in R4^-/-^ MEFs. To further confirm these findings, we investigated localization of β-catenin in R4^-/-^ MEFs to determine if they showed increased nuclear localization. First, our immunofluorescence images supported the 2-fold increase of total β-catenin levels as shown by western blot (Fig. 5A, B). They also showed significantly elevated β-catenin signals in the nucleus of R4^-/-^ MEFs, validating its higher activity in comparison to WT (Fig. 5B). Finally, as RHBDL4 participates in the proteasomal degradation of various substrates and β-catenin is degraded in the proteasome in the absence of Wnt signaling (48), we assessed if the absence of RHBDL4 induced impairment of proteasomal β-catenin degradation, which may ultimately lead to the stabilization of β-catenin. For this purpose, we treated WT and R4^-/-^ MEFs with the proteasomal inhibitor MG132. While as expected, β-catenin levels augmented with MG132 treatment in the WT MEFs, proteasomal inhibition did not affect β-catenin levels in the R4^-/-^ MEFs (Fig. 5C). In fact, β-catenin expression in WT cells under MG132 treatment mirrored β-catenin expression at basal condition in R4^-/-^ MEFs (Fig. 5C). This argues that dysfunctional proteasomal degradation is a likely cause for higher β-catenin levels in R4^-/-^ MEFs. Our conclusion is supported by our RNAseq data which showed no significant change in the expression of β-catenin (gene name *CTNB1*) at the transcriptional level.

**Figure 5:**
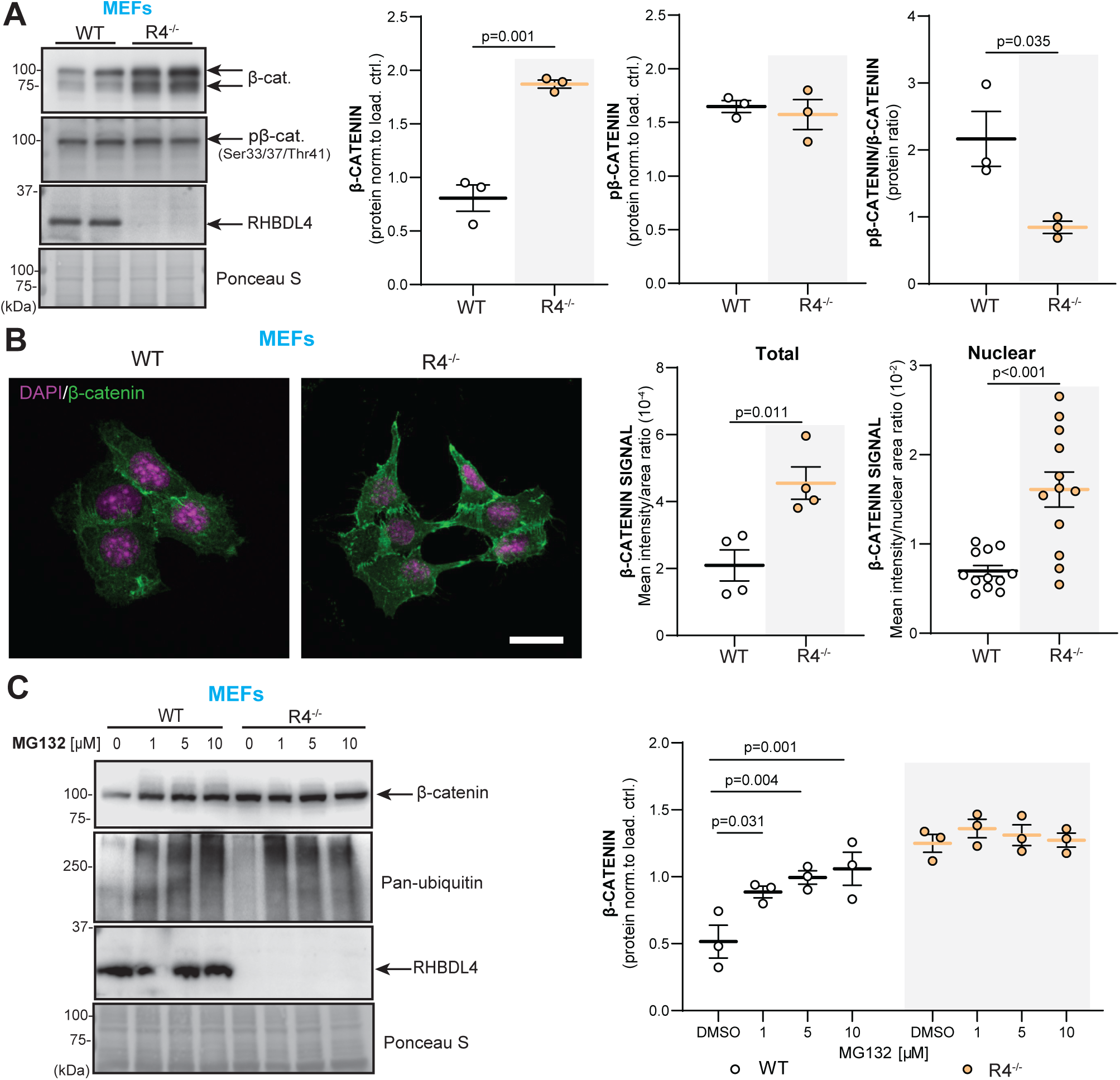
RHBDL4 null MEFs have upregulated β-catenin expression and activity due to decreased proteasomal degradation. A. Total β-catenin and pβ-catenin expression as well as pβ-catenin/β-catenin ratio in R4^-/-^ and WT MEFs. Expression was quantified and normalized to Ponceau-S. n= 3, mean ± SEM; Two-tailed, unpaired t-test, significant p-values reported. B. Total and nuclear β-catenin abundance in R4^-/-^ and WT MEFs. The nucleus is stained with DAPI (purple) while β-catenin is in green in the merged images. Confocal images taken at different z-planes were summed (step size 0.22 μm). Mean signal intensity per image normalized to cell area was quantified for total β-catenin expression while mean signal intensity normalized per nuclear area was quantified for nuclear β-catenin expression. Representative images of two biological replicates shown, (4 images quantified per genotype, 3 nuclei quantified per image) mean ± SEM; Two-tailed, unpaired t-test, significant p-values reported. C. Total β-catenin expression in R4^-/-^ and WT MEFs after MG132 treatment. Pan-ubiquitin is used as a control for successful proteasomal inhibition. Expression was quantified and normalized to Ponceau-S. n= 3, mean ± SEM; Two-way ANOVA (p=0.029 for the interaction, p=0.009 and p<0.001 for the treatment main effect) with Tukey’s multiple comparison test, significant p-values for post hoc analysis reported.

### Female APPtg mouse brains show increased **β**-catenin expression and activity in the absence of RHBDL4

Building on our *in vitro* findings, we sought to establish whether β-catenin expression was affected in APPtg mice in the absence of RHBDL4. First, when comparing WT and R4^-/-^ brains for both females and males, total β-catenin expression was increased similarly to what we observed in MEFs (Fig. 6A, B). However, pβ-catenin levels were also higher in both male and female R4^-/-^ mice than in WT (Fig. 6A, C), leading to no significant difference in pβ-catenin/β- catenin ratio between R4^-/-^ and WT brains (Fig. 6D). This discrepancy between the findings in MEFs and whole brain lysates may be attributed to the various cell types in the brain expressing RHBDL4 at different levels, which we do not resolve here. Intriguingly, when comparing APPtg mice to the other groups, we found a sex-specific trend that mirrored our results from the Y maze and NOR tests. For females, APPtg mice had comparable total β-catenin levels to WT but 2-fold increased pβ-catenin, suggesting low β-catenin activity in APPtg brains as shown by their high pβ-catenin/β-catenin ratio (Fig. 6A-C). Ablation of one or both alleles of RHBDL4 in APPtg females led to higher total β-catenin levels while pβ-catenin levels remained comparable, thus improving the pβ-catenin/β-catenin ratio (Fig. 6A-D). For males, APPtg mice had higher total β-catenin and pβ-catenin levels than WT, thus maintaining similar β-catenin activity as suggested by their similar pβ-catenin/β-catenin ratios (Fig. 6A-D). No significant difference in pβ-catenin/β-catenin ratio was observed for the other groups (Fig. 6 D), supporting our findings in Y maze and NOR tests where we conclude that the males are not impaired (Fig. 3). We conclude that the increase in β-catenin expression and activity is a likely cause for the cognition rescue we have observed in those groups (Fig. 3). In female mice, the ratio correlates with cognition, while male mice may have been too young to show impairment consistent with the β-catenin activity.

**Figure 6:**
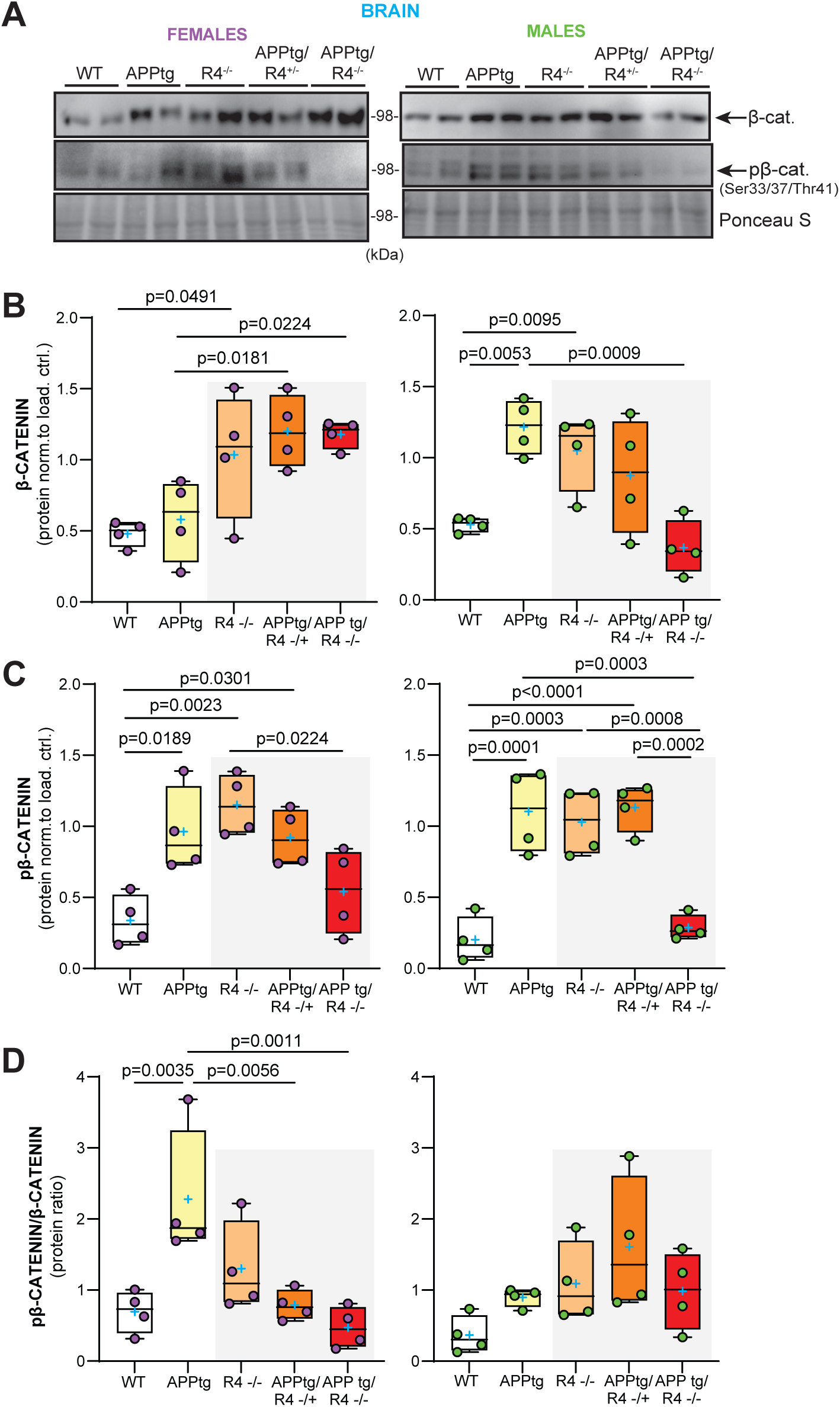
RHBDL4 knockout heterozygosity or homozygosity in APPtg mice normalizes β-catenin activity in APPtg females. A-C. Total β-catenin and pβ-catenin expression in brain lysates of female (purple) and male (green) WT, APPtg, R4^-/-^, APPtg/R4^-/+^, and APPtg/R4^-/-^ mice. Expression was quantified and normalized to ponceau-S. n= 4 per group, Box and whisker plots represent minimum to maximum values with median center lines while blue “+” represents the mean. Two-tailed, unpaired t-test performed between WT and R4^-/-^ group and one-way ANOVA (p=0.003 for females and p=0.001 for males) with Dunnett’s multiple comparison test where APPtg is compared to all other groups for β-catenin graphs. One-way ANOVA (p=0.002 for females and p<0.001 for males) with Tukey’s multiple comparison test for pβ-catenin graphs, significant p-values for post hoc analysis reported. D. pβ-catenin/β-catenin ratio in brain lysates of female (purple) and male (green) WT, APPtg, R4^-/-^, APPtg/R4^-/+^, and APPtg/R4^-/-^ mice. n= 4 per group, Box and whisker plots represent minimum to maximum values with median center lines while blue “+” represents the mean. one-way ANOVA (p=0.002 for females and p=0.093 for males) with Dunnett’s multiple comparison test for female graph, where APPtg is compared to all other groups. significant p-values for post hoc analysis reported.

## Discussion

We have previously described RHBDL4 as a regulator of APP biology in cell culture. We showed that RHBDL4 was a major producer of physiologically relevant A_η_-like peptides and competed with other significant APP processing pathways such as the amyloidogenic pathway (21, 23). Herein, we demonstrate the relevance of RHBDL4 in APP homeostasis *in vivo* and more importantly describe a new role of RHBDL4 in AD through its β-catenin regulating function. For APP regulation, we found that total APP levels were increased in the absence of RHBDL4. This due to the lack of RHBDL4-mediated APP degradation, allowing for an increase in β-CTF and Aβ production, in particular Aβ_40_. These findings corroborate our previous work in cell culture and solidify RHBDL4 as a regulator of APP levels in the brain (Fig. 7). In the context of AD, we found that RHBDL4 expression was increased in AD patients as well as in an AD mouse model. We speculate that RHBDL4 expression is upregulated as a response to ER stress and maybe intended to degrade excess APP (30, 32). Surprisingly, the increased expression of RHBDL4 was correlated with poorer cognition outcomes in AD patients. We propose that sustained RHBDL4 upregulation may lead to unwanted deleterious effects such as downregulation of the Wnt/β-catenin signalling pathway. This is in accordance with the fact that unresolved ER stress leads to maladaptive UPR, promoting apoptosis and inflammation instead of protective pathways (32). Our follow-up results in APPtg mice are consistent with the human data since cognitively impaired APPtg mice with decreased or absent RHBDL4 expression showed maintained memory. It is important to state that in models with rescued cognition, we also found increased Aβ production, which suggests that memory maintenance does not correlate with Aβ levels. Such findings are supported by previous studies where treatments maintaining cognitive performance in APPtg model did not have a lowering effect on Aβ production (49–51). The lack of correlation of cognition with Aβ levels underlines the importance of further developing non-Aβ targeting therapeutic strategies.

**Figure 7:**
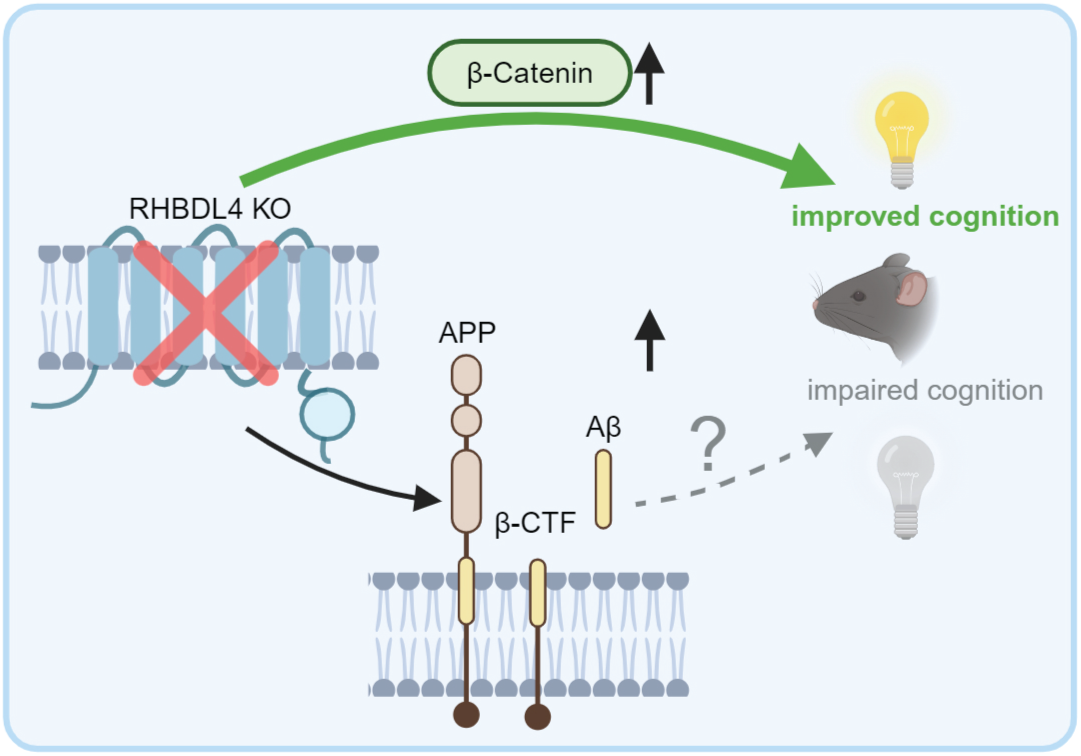
RHBDL4 in the context of an APP transgenic AD model. RHBDL4 ablation in an AD model increases APP expression and the production of amyloidogenic processing markers (β-CTFs and Aβ40). This confirms the relevance of RHBDL4 as a physiologically relevant modulator of APP. However, early in the pathology, β-catenin activity is normalized, and cognition is maintained in the absence of RHBDL4, despite the expected deleterious effects of the Aβ accumulation on cognition. These findings are in concordance with increased RHBDL4 expression in late-stage AD patients (Fig. 1), which negatively correlates with cognition. Thus, RHBDL4 may influence cognition by balancing the amounts of APP processing versus β-catenin signalling.

Using RNAseq data from R4^-/-^ MEFs and targeted approaches in our mouse samples, we posited β-catenin upregulation as the mechanism for memory rescue in the AD mouse model and put forward RHBDL4 as a novel negative regulator of β-catenin (Fig. 7). Decreased Wnt/β-catenin signalling has been implicated as an active player in AD for decades (42, 43, 52). The discovery of RHBDL4 as a modulator of this pathway is crucial and opens novel therapeutic avenues for AD. While we showed here that the increase in β-catenin in R4^-/-^ MEFs is likely due to improper proteasomal degradation, it is still unclear if RHBDL4 affects β-catenin directly or indirectly. On the one hand, RHDBL4 could directly bind to ubiquitinated β-catenin via its UIM and target it for proteasomal degradation. As RHBDL4 is in part located in the peri-nuclear areas of the ER (30) and our BioD data show that RHBDL4 interacts with nuclear pore complex components like nucleoporin NDC1 and integral proteins of the inner nuclear membrane such as MAN1 (29), it is possible that RHBDL4 encounters β-catenin while it translocates to the nucleus. Our BioID data also revealed α-catenin as a putative interactor of RHBDL4 (29). Interestingly, α-catenin is a known negative regulator of the Wnt/β-catenin signalling pathway via direct cytosolic sequestering of β-catenin or interaction with adenomatous polyposis coli (APC), a member of the destruction complex responsible for β-catenin degradation. We could thus speculate that RHBDL4 and α-catenin act together to regulate β-catenin degradation. On the other hand, our RNAseq data revealed that LRP6 and the Frizzled family of proteins are upregulated in R4^-/-^ MEFs. These proteins are crucial positive upstream regulators of β-catenin (39), such that their upregulation could lead to β-catenin stabilization. As they are transmembrane proteins that are trafficked through the ER, they could be substrates for RHBDL4. This is supported by our observation that LRP6 protein levels are also greatly increased (Fig. 4D). However, no substrate discovery study has identified them as substrates yet (31).

It is important to mention that a few studies have previously implicated RHBDL4 in Wnt/β-catenin signalling in the context of cancer (53, 54). In particular, Zhang et al. showed that in the HCT-116 cancer cell line, RHBDL4 knockdown led to decreased total β-catenin and decreased β-catenin phosphorylation at inhibitory positions serine 33, serine 37 and threonine 41. They further showed a positive correlation between RHBDL4 expression in tumor tissue and Zeb-1 expression, a downstream target of Wnt/β-catenin. However, they did not report directly on β-catenin and pβ-catenin expression in tumor tissue. While those results are overall contradictory to the ones described in this present report, the physiological contexts of the results may serve to explain the discrepancy. Our results were collected in physiologically normal cell lines and brain tissue as well as in APPtg mice. Zhang et al. studied the effects of RHBDL4 knockdown solely in the context of cancer cells lines or colorectal tumour tissues. Interestingly, it is known that AD and cancer are inversely correlated (55), which further suggests the importance of the disease *vs.* non-diseased context when assessing protein function. Taken together, we propose that RHBDL4 is an important regulator for Wnt/β-catenin signaling but that its effects are dependent on the physiological environment.

Another compelling finding of our study is the sex-specificity of the cognitive defects and rescue. We found that at 5 months of age, only female APPtg mice showed memory deficits while males performed similarly to controls. These findings are not surprising as it is known that women are disproportionately more affected by AD than men and female AD mouse models exhibit worse AD pathology than their male counterparts (56, 57). We further showed that female APPtg mice had high pβ-catenin/β-catenin as compared to WT, which was rescued by RHBDL4 knockout or heterozygosity. APPtg males on the other hand did not show a significant difference in their pβ-catenin/β-catenin ratio as compared to WT which supports the hypothesis that the memory rescue in females was due to increased β-catenin activity. Of note, testosterone treatments have been described to potently upregulate β-catenin expression in androgen receptor expressing tumors (58) and promote β-catenin translocation and transcriptional activity in 3T3-L1 preadipocytes (59). Furthermore, it is known that testosterone levels decrease with age in mice (60). This potentially explains why male APPtg mice are not impaired at 5 months of age and still have adequate β-catenin expression and activity. A follow-up study with older mice could help confirm this hypothesis. Such a study could determine whether the beneficial effects of RHBDL4 absence on cognition in females are maintained with time.

Overall, we propose that in APPtg mice and in AD, RHBDL4 is upregulated to control APP expression. However, RHBDL4’s upregulation also may have deleterious effects on the degradation of other substrates it regulates. In this case, β-catenin expression and activity are negatively affected by RHBDL4, likely promoting neurodegeneration and cognitive dysfunction (Fig. 7).

## Methods

### Cell culture and treatment

Immortalized embryonic fibroblasts from wild type and Rhbdl4-deficient mice (kindly provided by Dr. Matthew Freeman, Oxford University) were cultured in “full” Dulbeccós modified Eagle medium (DMEM) containing 4.5 g/l glucose, 0.584 g/l L-glutamine and 0.11 g/l sodium pyruvate (Wisent) and supplemented with 10% FBS (Wisent), at 37°C and 5% CO2. 1.5 x 10^5^ cells/well were seeded in “full” DMEM supplemented with 10% FBS in a 12-well plate for 48 h for adequate cell proliferation. For proteasomal inhibition experiments, 3 h treatment with 0-10 μM MG132 (Millipore) or DMSO, as vehicle control, was performed. 48 h post seeding, cells were lysed with TNE-lysis buffer (50mM Tris, pH 7.4, 150 mM NaCl, 2 mM EDTA, 1% NP40, and cOmplete protease inhibitors, Roche) and prepared for SDS-polyacrylamide gel electrophoresis (SDS-PAGE). 6 x SDS sample buffer (2 M Tris/HCl pH 6.8, 20% SDS, 100% glycerol, bromophenol blue, 500 mM DTT) was added to the samples for a final concentration of 1 x.

### Immunocytochemistry

5 x 10^4^ cells were seeded on coverslips in DMEM, supplemented with 10% FBS. 4% ice-cold Paraformaldehyde (PFA, Fisher) was used to fix cells for 15 mins at room temperature (RT) followed by 3 PBS washes. PFA was subsequently quenched with 500 mM NH_4_Cl for 10 min at RT and then cells were permeabilized with 0.1% Triton-X-100 (Sigma) for 10 mins at RT. Following this, cells were washed 5x with PBS, blocked with 3% normal goat serum (NGS, Sigma) for 1 h at RT, and finally incubated with 1:200 dilution of mouse anti-β-catenin (Cell signaling, L52E2 2677S) in blocking buffer overnight at 4°C. The next day, 3 PBS washes were carried out for 5 mins at RT before incubation with Alexa 488-coupled secondary antibodies directed against rabbit IgG (Life Technologies) diluted at 1:2000, RT in the dark for 1h. After washing off the secondary antibody with PBS, the coverslips were mounted on microscope slides using fluoroshield with DAPI (Sigma). Confocal images were acquired by Zeiss LSM 800 confocal laser scanning microscope and analyzed using ImageJ.

### MEFs RNAseq

5 x 10^5^ cells/well were seeded in “full” DMEM supplemented with 10% FBS in a 6-well plate for 48 h for adequate cell proliferation. The cell medium was aspirated, and cells were washed in PBS once before being harvested in fresh PBS. Cells were spun down at 1500 rpm for 3 minutes. The supernatant was aspirated, and the cell pellet was flash-frozen. The QIAGEN RNeasy Micro Kit was utilized for RNA extraction, and the quality of the extracted RNA was verified by Bioanalyzer. mRNA enrichment was carried out on 500 ng total RNA using the NEBNext Poly(A) mRNA Magnetic Isolation Module Kit, followed by library generation using the Swift RNA Library Kit. Subsequent sequencing occurred on either an Illumina NovaSeq instrument at the La Jolla Institute for Allergy and Immunology Sequencing Core or a HiSeq 4000 at the Michael Smith Genome Sciences Centre in Vancouver, Canada. To ensure result consistency, three samples were sequenced at both locations. Alignment to hg19 was performed with the STAR aligner (v2.5.3a) using default parameters on FASTQ files. RNA-SeQC analyzed the resulting Bam files to confirm library quality. Read counts were computed with htseq-count, and RPKM values were determined using cufflinks (v2.2.1) with default settings. Principal component analysis (PCA) was conducted using the prcomp function in R and visualized with the ggfortify package. Genes with RPKM values below 2 in all samples were excluded from subsequent analysis. Differential expression between RHBDL4 KO and WT MEFs was computed from transcript counts for each gene using the DEseq2 software package. Log2FC and the FDR-adjusted p-value were used for downstream analysis. The threshold for differential expression was set to 1.8-fold change and statistical significance was set at p-value < 0.05. EnrichR was used for pathway analysis. MATLAB (v23.2.0.2428915) was employed to visualize and filter the gene expression patterns across biological replicates.

### Protein extraction from mouse tissue

For the analysis of protein expression in mouse, similar frontal cortical brain regions, liver lobes, and pancreas pieces were snap frozen after dissection and stored at −80°C until further analysis. For tissue lysis, 5 times the volume of tissue homogenization buffer (20 mM Hepes, 150 mM NaCl, 20% glycerol, 2 mM EDTA, 1% NP-40, 0.1 % sodium deoxycholate, pH 7.4) supplemented with 2 x cOmplete Protease Inhibitors was added to 100-200 mg of tissue. After mechanical tissue homogenization, lysates were incubated at 4°C for 1 h and then spun down at 20 000 g for 1 h, 4°C. Lysates were adjusted to equal protein concentrations (3 μg/μl) and prepared for SDS-PAGE. 6 x SDS sample buffer was added to the samples for a final concentration of 1 x.

For analysis of human Aβ from APPtg mice, 50 mg posterior cortical regions were homogenized with 5 times the volume of tissue homogenization buffer containing 2 x cOmplete Protease Inhibitors. Following mechanical homogenization, homogenates were spun down at 5000 g for 10 minutes, 4°C. The supernatant was collected and diethylamine DEA buffer (0.4% DEA, 0.1 M NaCl) added at 1:1 ratio. The DEA-homogenate mixture was spun down using an ultracentrifuge at 120 000 g for 1h, 4°C. The resulting supernatant was neutralized with 0.5 M Tris-HCL (pH 6.8) and was used as the soluble Aβ fraction. The remaining pellet after centrifugation was dissolved with 70% formic acid and the solution was spun down at 120 000 g for 1h, 4 °C. The supernatant was collected and neutralized with formic-acid neutralization buffer (1M Tris base, 0.5 M Na_2_HPO_4_, 0.05% NaN_3_), yielding the Aβ insoluble fraction. A total of volume of 25 μl per sample was loaded on the Meso Scale Discovery (MSD) immunogenicity assay triplex plate (6E10). Aβ38, Aβ40, and Aβ42 levels were quantified according to the manufacturer’s instructions.

### Western blot

Samples were separated on 4-12% bis-tris gels (Novex, Nupage, Invitrogen) or 10% and 15% tris-glycine. Bis-tris gels were run with MES running buffer (Invitrogen). Proteins were transferred onto nitrocellulose (chemiluminescence imaging) or to Immobilion FL PVDF membranes (Millipore, fluorescence imaging) using transfer buffer with 10% ethanol. Western blots were blocked with 5% skim milk in TBS-T (chemiluminescence) or with Odyssey Blocking buffer (Li-COR Biosciences, fluorescence). The following primary antibodies were used: rabbit-anti-RHBDL4 (HPA013972 Sigma), rabbit anti-β-Tubulin (D2N5G, Cell Signaling), Y188 (APP C-terminus, ab32136, Abcam), mouse-anti-β-actin (8H10D10, Cell Signaling), rabbit anti-LRP6 (C47E12, Cell Signaling), rabbit anti-β-catenin (9562S, Cell Signaling), rabbit anti-Phospho-β-Catenin Ser33/37/Thr41 (Cell Signaling, 9561S), mouse anti-ubiquitin (Cell signalling, P4DI 3936S). Horseradish peroxidase (HRP)-coupled secondary antibodies directed against mouse or rabbit IgG were purchased from Promega for chemiluminescence imaging. Fluorophore-conjugated antibodies IRDye 800 anti-rabbit (Li-COR Biosciences) was used for fluorescence imaging. Chemiluminescence images were acquired using the ImageQuant LAS 500 or 600 system (GE Healthcare) and analyzed using imageJ. Fluorescence images were obtained with the Odyssey CL-X imaging system, processed with Image Studio Lite V4.0. and analyzed using imageJ. GAPDH or ponceau-S signals were used for normalization of proteins of interest expression.

### mRNA extraction & qPCR analysis from mouse tissue

mRNA from 50 mg of similar frontal cortical brain regions was extracted using TRIzol Reagent following the manufacturer’s protocol (Invitrogen) and then reverse transcribed into cDNA (iScript mix). qPCR was performed using the SsoAdvanced SYBR green supermix on a Biorad CFX384Touch cycler. Primers used were: RHBDL4 forward: (CCTTCAGGACAGCAGAACCACT) and reverse (GCTGTGTAGACATCATAGTTCCTG), RSP18 forward (CGGAAAATAGCCTTCGCCATCAC) and reverse (ATCACTCGCTCCACCTCATCCT), GAPDH forward (CATCACTGCCACCCAGAAGACTG) and reverse (ATGCCAGTGAGCTTCCCGTTCAG). Primer efficiency and optimal melting temperature were determined to be 90-110% and 57°C, respectively. For normalization of *Rhbdl4* gene expression, RSPI8 and GAPDH were confirmed to be stable and thus were used as reference genes. RT-qPCR was analyzed using the CFX Maestro software. Data are displayed as mean expression.

### Y maze

5-month-old animals were placed in a Y-Maze with the dimensions of 40 cm x 8 cm x 10 cm with the arms at an angle of 120°. The animal was allowed to explore the maze freely for 5 minutes while being recorded. An arm entry was recorded when all four limbs of the animal wholly entered the arm. Over multiple entries, cognitively healthy animals show a preference to enter a less recently visited arm. A full alternation is counted when the mouse visits 3 different arms in a row. The experiments were performed in a dark room with only red light and recorded using an infrared camera. Spontaneous alternating performance (SAP) is reported, where it is calculated as: (number of alternations/number of entries-2) x 100.

### Novel object recognition

5-months-old animals were first “habituated” to an empty open field box in a dark room (acclimatation). 24 h later, mice were exposed to two identical, non-stress inducing “familiar” objects (familiarization) placed in two opposing corners of the box. After a retention interval of 24 h, the animals were returned to the box in which one of the two objects was exchanged with a novel object (testing). Each session lasted 5 min, during which mice were allowed to interact with the objects freely and were recorded using an infrared camera. All objects were randomized between animals. Analysis of the time spent with each object was conducted using the ODlog software. Index of preference is reported, where it is calculated as: time with novel object / (time with familiar + time with novel) x 100.

### Mouse housing and cross

Homozygous RHBDL4-knockout (R4^-/-^) mice are viable and show no obvious phenotypes (30). APP transgenic J20 (APPtg) mice are a well described AD mouse model (35). Mice are housed in the Munter lab colony according to the McGill University standard operating procedure mouse breeding colony management #608. All procedures were approved by McGill’s Animal Care Committee and performed in accordance with the ARRIVE guidelines (Animal Research: Reporting in Vivo Experiments). R4^-/-^ mice were bred with APPtg mice to obtain APPtg/R4^-/+^ mice. APPtg/R4^-/+^ x APPtg/R4^-/+^ breeders were then set up to obtain WT, APPtg, R4KO, APPtg/R4^-/+^ and APPtg/R4^-/-^. Genotyping was performed by Transnetyx.

### Human brain sample analysis

Frozen post-mortem dorsolateral prefrontal cortex came from participants in the Religious Orders Study or Rush Memory and Aging Project (ROSMAP) was performed as described (61). Detailed methods of both cohort studies have been previously reported (62, 63). Both studies were approved by the Institutional Review Board of Rush University Medical Center. Each participant signed an informed consent, Anatomic Gift Act, and repository consent to allow their data to be repurposed. A total of 21 cognitive performance tests were used in both studies. The clinical diagnosis of Alzheimer’s dementia was uniform and structured and followed the NINCDS-ADRD criteria as reported (64). Nineteen of the tests were z-scored and averaged for a global measure of cognition (65). The pathologic diagnosis of AD was made by NIA-Reagan criteria as previously described (34). A global measure of AD pathology, β-amyloid load, PHFtau tangle density, cortical Lewy bodies, other neurodegenerative diseases, and macro- and micro-vascular disease were assessed as previously described (66–69). mRNA expression from frozen post-mortem dorsolateral prefrontal cortex extraction was performed from the gray matter of the dorsal lateral prefrontal cortex, followed by next-generation RNA sequencing (RNA-Seq), which was performed on the Illumina HiSeq (RNA integrity score > 5; quantity threshold > 5 µg), as previously described (70, 71). Fragments were quantile-normalized per kilobase of transcript per million fragments mapped (FPKM).

We used adjusted FPKM values for analysis. *RHBDL4* mRNA expression was determined for cases with pathologic AD (n=321) and those without pathologic AD. Level of transcript expression between pathologic AD and no pathologic AD were first compared using two-sample t-test. To test further the association of *RHBDL4* mRNA expression with a pathologic diagnosis of AD (yes vs. no) and cortical Lewy bodies (yes vs. no), we ran logistic regression, controlling for age at death, sex and education. To assess the association with levels of AD pathology, β-amyloid, neurofibrillary tangles, macro- and micro-infarcts, we ran linear regressions with the pathologies being the dependent variables, controlling for age at death, sex and education. Level of AD pathology was log transformed, and β-amyloid was square-root transformed to improve normality. Finally, we used linear regression to test the association with global cognition. We used Box-Cox with negatives, a modified version of the Box-Cox transformation that allows for negatives responses, to transform global cognition to improve its normality.

For protein studies, human post-mortem parietal cortex tissues from AD and controls were from the University of California, Irvine, Center for Brain Aging & Dementia Tissue Repository. Tissue homogenates were prepared as previously described (72). 50 ug of each homogenate was mixed with Laemmli sample buffer (0.0625 M Tris-HCl pH 6.8, 2% SDS, 10% glycerol, 5% 2mercaptoethanol, 0.001% bromophenol blue) and incubated overnight at 4 C. Samples, without heating, were loaded onto and resolved on 12% SDS-PAGE gels. After transferring the proteins to PVDF membranes (Millipore, Etobicoke, ON, Canada) as described, the membranes were blocked with PBS mixed 1:1 with Odyssey Blocking Buffer (LI-COR Biosciences, Lincoln, Nebraska, USA), washed with double distilled water, dried and stored until probing with the antibodies. Lastly, protein lysates from additional ROSMAP brain samples were prepared following the protocol above. In brief, 250mg human brain tissue were homogenised in 50 mM Tris, 150 mM NaCl, 10 mM EDTA and 2x cOmplete protease inhibitor using the Miltenyi Gentle Max. 1% NP-40 and 0.1% deoxycholate were added after and samples were spun down at 15000g for 10 min. The protein concentration per sample was adjusted to 4 µg/µl. 4x LDS sample buffer was added to a final concentration of 1x and samples were incubated over night at 4°C. Western blot analysis was performed as mentioned above. RHBDL4 protein levels were quantified and normalised to β-actin+GAPDH. Results from both cohorts were combined and after a significant Shapiro Wilk test for normality, a Mann-Whitney U test was performed to assess significance.

### Statistical analysis

Statistical data analysis was performed with GraphPad Prism 10 unless described otherwise (human data). Details as indicated in the figure legends.

## Acknowledgements

We thank Dr. Scott De Vito for his help with confocal microscopy images. Image collection was performed in the McGill University Advanced BioImaging Facility (ABIF), RRID:SCR_017697. We thank Dr. Joseph Flores at the Lady Davis Institute for Medical Research for his help collecting MSD ELISA data. We thank Dr. Judith Mandl, McGill University, for her advice and contribution to figure conception. The RHBDL4 KO mouse model as well as WT and RHBDL4 KO mouse embryonic fibroblasts were kindly provided by Dr. Matthew Freeman, University of Oxford. The McGill-Thy1-APP mouse strain was kindly provided by Dr. Claudio Cuello, McGill University. This research was supported by NSERC Discovery grant no. RGPIN-2015-04645, the Canadian Institutes of Health Research (CIHR) PJT-175306 and PJT-162302, Canada Foundation of Innovation Leaders Opportunity Fund (CFI-LOF, 32565), Alzheimer Society of Canada Research Grant 20-02, Fonds d’innovation Pfizer-FRQS sur la maladie d’Alzheimer et les maladies apparentées no. 31288, and The Scottish Rite Charitable Foundation of Canada 16112. Support was received from CIHR (PJT-159493) to MF. ROSMAP is supported by P30AG10161, P30AG72975, R01AG15819, R01AG17917. U01AG46152, and U01AG61356. ROSMAP resources can be requested at https://www.radc.rush.edu and www.synpase.org. YP received the Davis fellowship through McGill’s Faculty of Medicine and a stipend from the Canada First Research Excellence Fund, awarded to McGill University for the Healthy Brains for Healthy Lives initiative. SJR received a CGS-Master’s award.

## Author contributions

YP designed and conducted experiments, analyses and manuscript preparation. SP analyzed RHBDL4 protein expression in AD patients brain samples. JY performed the RNAseq analysis from AD patients’ brain and completed correlation studies. SJR analyzed RHBDL4 expression in various mouse tissues, conducted behavior test in McGill-Thy1-APP and collected R4^-/-^ MEFs samples for RNAseq analysis. MR, BX, JS, HMM and AN participated at various stages of RNAseq analysis for the R4^-/-^ MEFs samples. BM produced the membranes with human brain lysates. MF supervised human brain lysate experiments. DB supervised analyses of AD patient RNAseq data. WP supervised R4^-/-^ MEFs RNAseq data. LMM supervised and coordinated the project, revised manuscript. All authors revised and approved the manuscript.

## Figure Legends

**Supplemental Figure 1:**
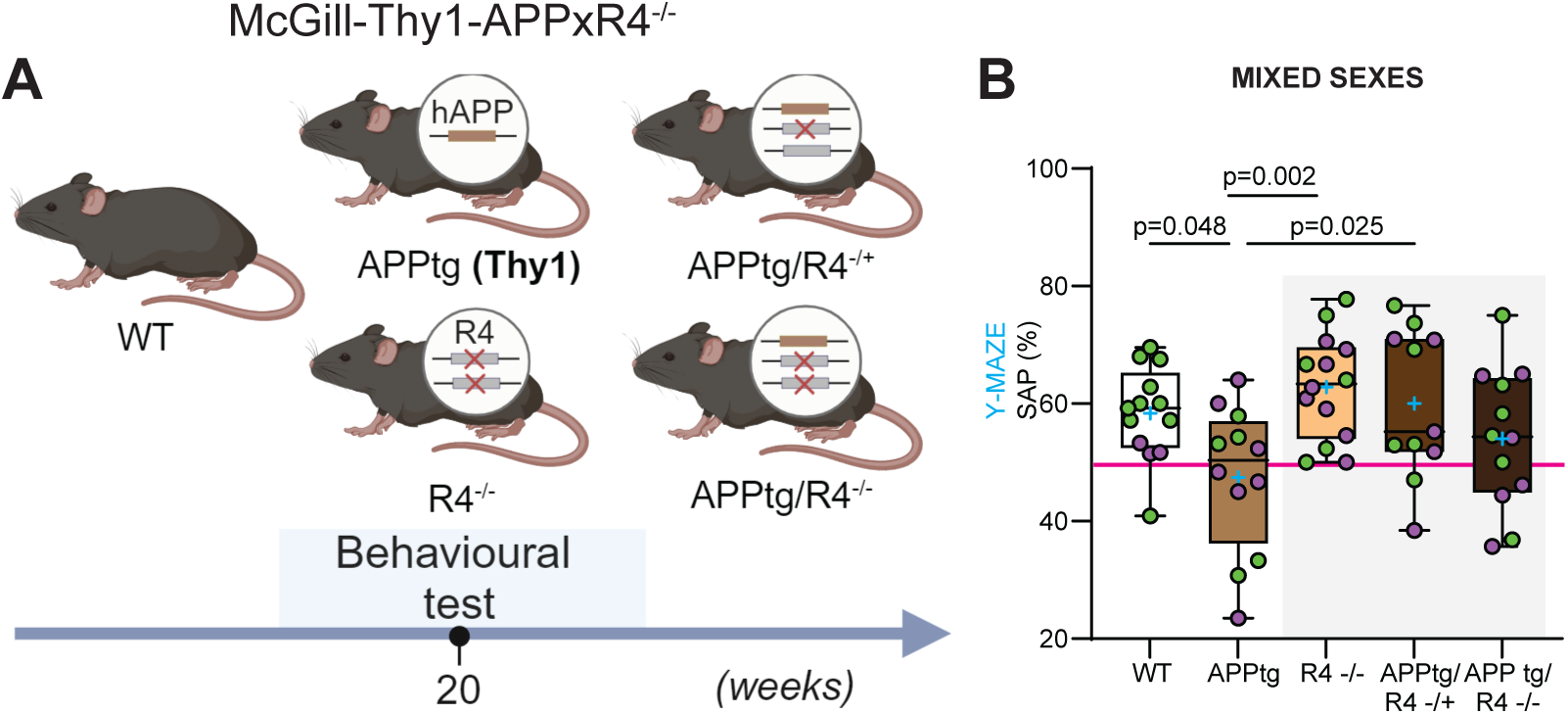
Cognitive defects are improved in the absence of RHBDL4 expression in a different APPtg model. A. Schematic representation of experimental design and analysis timeline for preliminary cohort of APPtg McGill-Thy1-APP mice crossed to the R4^-/-^ model. B. Spontaneous alternation performance (SAP) score from Y maze test of WT, APPtg, R4^-/-^, APPtg/R4^-/+^, and APPtg/R4^-/-^ mice. Female data points are in purple and male in green, n=12-14 per group. Box and whisker plots represent minimum to maximum values with median center lines while blue “+” represents the mean. One-way ANOVA (p=0.008) with Dunnett’s multiple comparison test, significant p-values for post hoc analysis reported.

